# ADGRG1 blockade drives astrocyte positioning into the extracellular matrix to reduce scar size

**DOI:** 10.64898/2025.12.18.695114

**Authors:** Gentaro Ono, Kazu Kobayakawa, Hirokazu Saiwai, Kazuya Yokota, Tetsuya Tamaru, Kazuhiro Hata, Hirotaka Iura, Yohei Haruta, Kazuki Kitade, Jun Kishikawa, Kiyoshi Tarukado, Kenichi Kawaguchi, Dai-Jiro Konno, Kinichi Nakashima, Takeshi Maeda, Seiji Okada, Yasuharu Nakashima

**Affiliations:** Department of Orthopaedic Surgery, Graduate School of Medical Sciences, Kyushu University, Fukuoka, Japan; Department of Energy and Materials, Faculty of Science and Engineering, Kindai University, Osaka, Japan; Department of Stem Cell Biology and Medicine, Graduate School of Medical Sciences, Kyushu University, Fukuoka, Japan; Department of Orthopaedic Surgery, Spinal Injuries Center, Iizuka, Japan; Department of Orthopaedic Surgery, Graduate School of Medicine, Osaka University, Suita, Japan

**Keywords:** ADGRG1, Astrocyte, Glial scar, Spinal cord injury, Type III collagen

## Abstract

After the central nervous system (CNS) injury, inflammatory cells enter the lesion core and a fibrotic extracellular matrix accumulates. Astrocytes subsequently cluster around this matrix, forming a glial scar, but the role of the extracellular matrix in scar formation remains poorly understood. Here, we identify type III collagen, a major fibrotic extracellular matrix component after spinal cord injury, as a key regulator of astrocyte behavior. Using single-nucleus transcriptomics, spatial transcriptomics, and time-lapse cell imaging, we show that type III collagen positions astrocytes at the boundary of the fibrotic matrix by activating the adhesion G protein-coupled receptor G1 (ADGRG1) and its downstream effector RhoA. Blocking ADGRG1 genetically or pharmacologically allows astrocytes to enter the fibrotic matrix, resulting in a smaller scar, improved neuronal regeneration, and better motor recovery. The migration of human-induced pluripotent stem cell-derived astrocytes was inhibited by type III collagen via the ADGRG1-RhoA pathway, supporting their therapeutic potential. Our findings reveal the importance of controlling glial scar positioning via the type III collagen-ADGRG1 axis, providing a novel therapeutic target for CNS injury.

## Introduction

Spinal cord injury (SCI) leads to devastating and permanent neurological deficits, mainly due to the formation of scars. These scars consist of infiltrating inflammatory cells, a dense layer of astrocytes—known as the glial scar—and a fibrotic extracellular matrix (ECM), which together create both physical and biochemical barriers that obstruct axonal regeneration (Anderson *et al*, 2016; Ayazi *et al*, 2022). The ECM is a major component of the scar and is known to influence a variety of cellular responses. For example, one type of ECM molecule, chondroitin sulfate proteoglycans (CSPGs), are well-known inhibitor of axonal growth and illustrate how ECM molecules can actively regulate cellular behavior after injury (Bradbury *et al*, 2002; Brown *et al*, 2012). More recently, type I collagen has been shown to promote astrocyte scarring (Hara *et al*, 2017), highlighting the influence of fibrotic ECM components on scar formation. Among these components, type III collagen is one of the most abundantly deposited fibrotic ECM proteins in the lesion core following SCI, and its gene expression is also upregulated after CNS injury (Hara *et al*, 2017; Wareham *et al*, 2024). However, in contrast to CSPGs and type I collagen, the role of type III collagen has been largely overlooked.

In this study, we identify type III collagen as a key regulator of astrocyte positioning during scar formation. We show that type III collagen activates the adhesion G protein-coupled receptor 1 (ADGRG1) and its downstream effector RHOA to entrap astrocytes at the edge of the fibrotic ECM. Genetic or pharmacological inhibition of this pathway enables astrocytes to enter the fibrotic ECM, leading to smaller glial scars, enhanced axonal regeneration, and improved motor recovery.

We uncover a previously unrecognized role of type III collagen as an organizer of CNS scar structure. Modulating the type III collagen–ADGRG1 axis may open new avenues for regenerative therapies in CNS injury.

## Results

### Role of type III collagen in astrocyte behavior

To investigate how astrocytes respond to spinal cord injury (SCI) over time, we analyzed publicly available single-nucleus RNA-sequencing (snRNA-seq) datasets of spinal cord tissue from uninjured mice and mice at multiple time points post-injury (Skinnider *et al*, 2024). Using unsupervised clustering and Cell Typist-based cell type annotation, we identified 49 distinct cellular clusters, including several astrocyte populations (Fig. 1A, Fig. S1A) (Domínguez Conde *et al*, 2022; Xu *et al*, 2023; Yao *et al*, 2023). Astrocytes were subdivided into five distinct clusters. Among them, Astro_1 represented astrocytes in the intact spinal cord, while Astro_3, 4, and 5 were specific to the acute phase post-injury. In contrast, Astro_2 was predominant during the subacute and later phases (Fig. 1B, Fig. S1B). To clarify how type III collagen, encoded by *Col3a1*, affects astrocytes, we examined pathways with CellPhoneDB (Efremova *et al*, 2020) Among the various *Col3a1*-associated pathways, the ADGRG1-mediated pathway exhibited the most potent interaction on astrocytes (Fig. 1C). While type III collagen affects several cell types—including astrocytes, endothelial cells, and ependymal cells—only the ADGRG1 pathway showed a substantial effect on astrocytes (Fig. 1D). Interestingly, this pathway displayed dynamic temporal regulation: its activity transiently declined during the hyperacute phase, gradually increased between days 4-7, and subsequently diminished again, suggesting this pathway may be one of the regulatory factors influencing glial scar formation by astrocytes in the subacute stage of SCI (Fig. 1E). Furthermore, spatial transcriptomic analysis revealed that Astro_1 localizes near cells strongly expressing *Col3a1*, particularly those belonging to the VLMC (vascular leptomeningeal cell) _1 cluster (Fig. 1F). These snRNA-Seq and spatial transcriptomic analysis supported that the type III collagen-ADGRG1 pathway dynamically regulates astrocyte behavior in the local microenvironment of the injured spinal cord.

**Figure 1.**
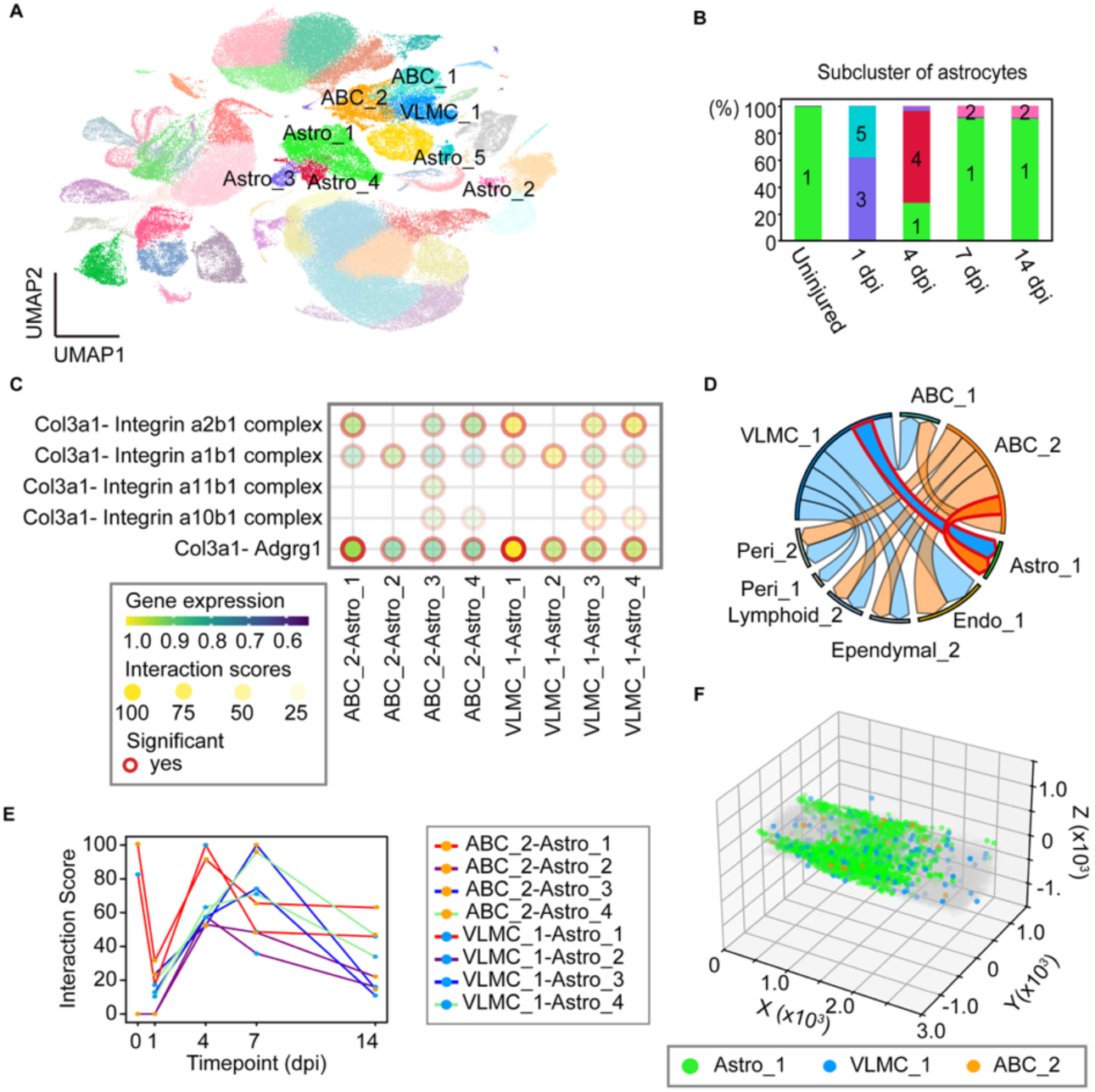
Type III collagen–mediated interactions affecting astrocytes after SCI. (A) UMAP of cell states (cell types) in the injured spinal cord snRNA-seq (n = 151712) datasets. Astro: Astrocytes, ABC: Arachnoid barrier cells, Endo: Endothelial cells, VLMC: vascular leptomeningeal cells. (B) Stacked bar plot showing the temporal changes in the relative proportions of astrocyte subclusters. dpi: days post-injury. (C) Dot plots showing the snRNA-seq analysis results concerning cell-cell interactions at 4 dpi. Color indicates co-expression of ligand and astrocyte receptor genes; transparency reflects the interaction score; presence of a red outline indicates statistical significance. P-values < 0.05 were considered significant. (D) Circos plots showing the number of significant and strong cell interactions via type III collagen in the injured spinal cord at 4 dpi. Numbers above the arrows indicate the number of pathways that were both significant and strong for each cell–cell pair. Arrows outlined in red indicate the COL3A1-ADGRG1 pathway. P-values < 0.05 were considered significant. Interaction scores > 70 were considered strong. (E) Line plots showing temporal changes in Type III collagen-ADGRG1 pathway interaction scores in the spinal cord. (F) 3D-spatial transcriptomics of the injured spinal cord at 7 dpi, highlighting type III collagen–secreting cells and neighboring astrocytes. The figure depicts the spatial localizations of Astro_1 (green), VLMC_1 (blue), and ABC_2 (orange).

### ADGRG1 inhibition enables astrocytes to enter the fibrotic ECM

We next investigated whether blocking the ADGRG1 pathway could change the structure of the glial scar in vivo. We first injected the ADGRG1 antagonist, dihydromunduletone (DHM), directly into the injured spinal cord (Stoveken *et al*, 2016a). Based on preliminary experiments with multiple administration methods (Fig. S2 A-E), we adopted a protocol involving a single injection at four days post-injury (dpi).

In DHM-treated mice, astrocytes were no longer confined to the edge of the fibrotic ECM but instead shifted their position into the fibrotic ECM, forming a smaller and more centrally located glial scar (Fig. 2 A–D). Furthermore, neuronal tracing with adeno-associated virus (AAV) -*Cag-tdTomato* injection into the sensorimotor cortex at 14 dpi revealed that the volume reduction of the glial scar induced by DHM injection led to extensive axonal regeneration (Fig. 2 E-F). However, the injection of DHM did not significantly affect the inflammatory response in the spinal cord, suggesting the promoted axonal regeneration by the antagonist of ADGRG1 was not via regulation of the inflammatory response (Fig. 2 H-I). The smaller glial scar volume resulted in axonal regeneration and consequently improved motor function (Fig. 2J). To verify that these effects were astrocyte-specific, we designed an AAV expressing a short hairpin RNA targeting *Adgrg1* (Adgrg1 K.D.) (Fig. S2F) and injected it intrathecally 7 days prior to SCI. Similar to pharmacological inhibition, genetic knockdown of *Adgrg1* led to astrocytes entering into the fibrotic ECM and formation of a smaller, more compact scar (Fig. 2 K-M). Notably, these structural changes were accompanied by improved motor function recovery compared to control AAV-treated animals (Fig. 2 N). These results indicate that the type III collagen–astrocytic ADGRG1 axis controls the spatial organization of astrocytes after SCI, and that inhibiting this pathway promotes beneficial remodeling of the glial scar.

**Figure 2.**
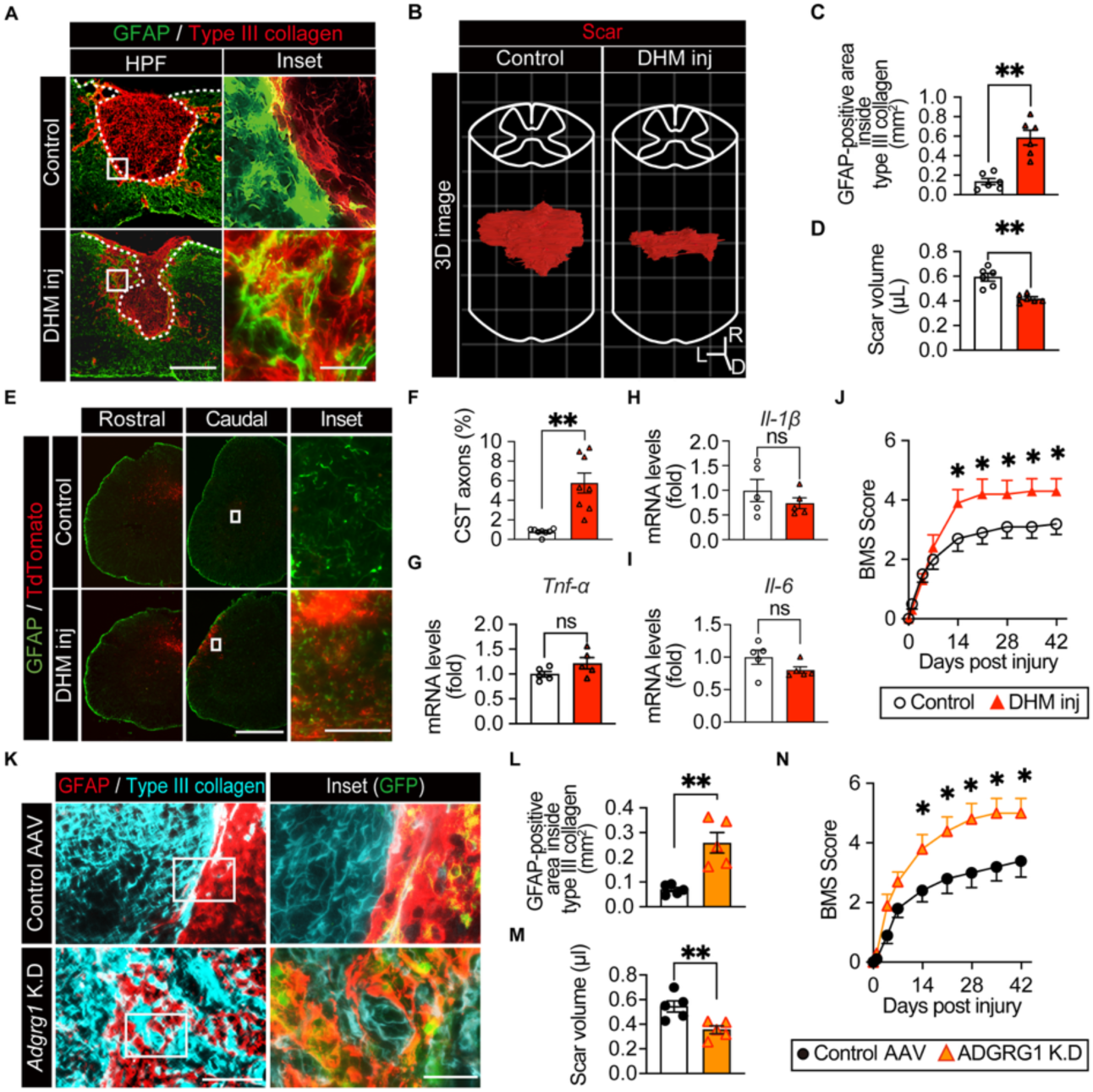
ADGRG1 inhibition reduces the volume of the glial scar and improves motor outcome after SCI. (A) Representative images showing GFAP (green) and type III collagen (red) staining of spinal cords at 14 dpi receiving PBS (control group) or DHM (DHM inj group). The white dotted line indicates the border of the glial scar. Scale bars: 500 μm and 50 μm (inset). (B) Three-dimensional images of the scar in the control group and in the DHM inj group (red: type III collagen-positive volume surrounded by GFAP-positive area). R indicates the rostral side, D indicates the dorsal side, and L indicates the left side. Grid size: 500 μm. (C) Quantitative analysis of the area of astrocytes inside type III collagen in the control group and in the DHM inj group (n = 6). (D) Quantitative analysis of scar volume between the control group and DHM inj group (n = 6). (E) Neuronal tracing images showing tdTomato (red)-positive axons and GFAP (green)-positive astrocytes in the control group and in the DHM inj group. Scale bars: 500 μm and 50 μm (inset). (F) Quantitative analysis of axons in the control group and in the DHM inj group (n = 8). (G-I) Changes in *Il-6*, *Il-1β*, and *Tnf-α* gene expression between the control group and the DHM inj group at 7 dpi (n = 5). (J) The time course of the BMS score after SCI in the control group and in the DHM inj group (n = 10). (K) Representative images showing GFAP (red), type III collagen (cyan), and GFP (green, inset) staining of spinal cords at 14 dpi, treated with control AAV (control AAV group) or AAV-*Gfap*-*shAdgrg1* (*Adgrg1* K.D group). Scale bars: 200 μm and 50 μm (inset). (L) Quantitative analysis of the area of astrocytes inside type III collagen in the control AAV group and in the *Adgrg1* K.D group (n = 5). (M) Quantitative analysis of scar volume between the control AAV group and the *Adgrg1* K.D group (n = 5). (N) The time course of the BMS score after SCI in the control AAV group and in the *Adgrg1* K.D group (n = 10). *P < 0.05, **P < 0.005, unpaired t-test (C, D, F-I, and L-M), two-way ANOVA (J and N). Data represent the mean ± s.e.m (C-D, F-J, and L-N).

### Mechanistic Insights into ADGRG1-Mediated Regulation of Astrocyte Migration

Given that ADGRG1 inhibition allowed astrocytes to enter the fibrotic ECM after SCI, we sought to elucidate the underlying mechanism using an in vitro model. Primary cultured astrocytes were first converted into a reactive phenotype and then plated under three different conditions: poly-L-lysine coating (control group), type III collagen coating (type III collagen group), and type III collagen coating with dihydromunduletone (DHM) treatment (DHM group) (Stoveken *et al*, 2016a). The migration ability of astrocytes was evaluated using a scratch assay. In the type III collagen group, we observed impaired astrocytic migration, while DHM treatment restored migration ability (Fig. 3 A and B). Detailed cell tracking revealed shorter process lengths in the type III collagen group (Fig. 3C, Fig. S3A). Cells migrate through the following stages: the protrusion of processes at the anterior edge of the cell, the adhesion of processes to the substrate, and cytoskeletal contraction (SenGupta *et al*, 2021). Therefore, the shorter the processes, the less distance the cell can migrate in a single motion. Conversely, in the DHM group, astrocytes exhibited a mixture of short and long processes. There was no significant difference in the distance of cell migration and mean speed between the DHM group and the control group, respectively (Fig. 3D, Fig. S3B). Since the type III collagen-ADGRG1 pathway is known to regulate cell migration via RHOA-GTP binding, we next evaluated RHOA-GTP binding levels in each group. The type III collagen group showed elevated levels of RHOA-GTP binding relative to the control group, but no significant differences were observed in RHOA-GTP binding levels between the DHM group and the control group (Fig. 3E). Furthermore, when we compared the effects of different collagen subtypes on astrocyte migration, we found that astrocytes cultured on type III collagen displayed significantly reduced migratory capacity compared to those on type I or type IV collagen (Fig. 3 F and G). These findings indicate that type III collagen activates the ADGRG1–RHOA pathway to position astrocytes along the edge of the fibrotic ECM after SCI, and that pathway inhibition enables astrocytes to migrate into the fibrotic ECM, which leads to an inward shift of the glial scar position, resulting in a smaller glial scar.

**Figure 3.**
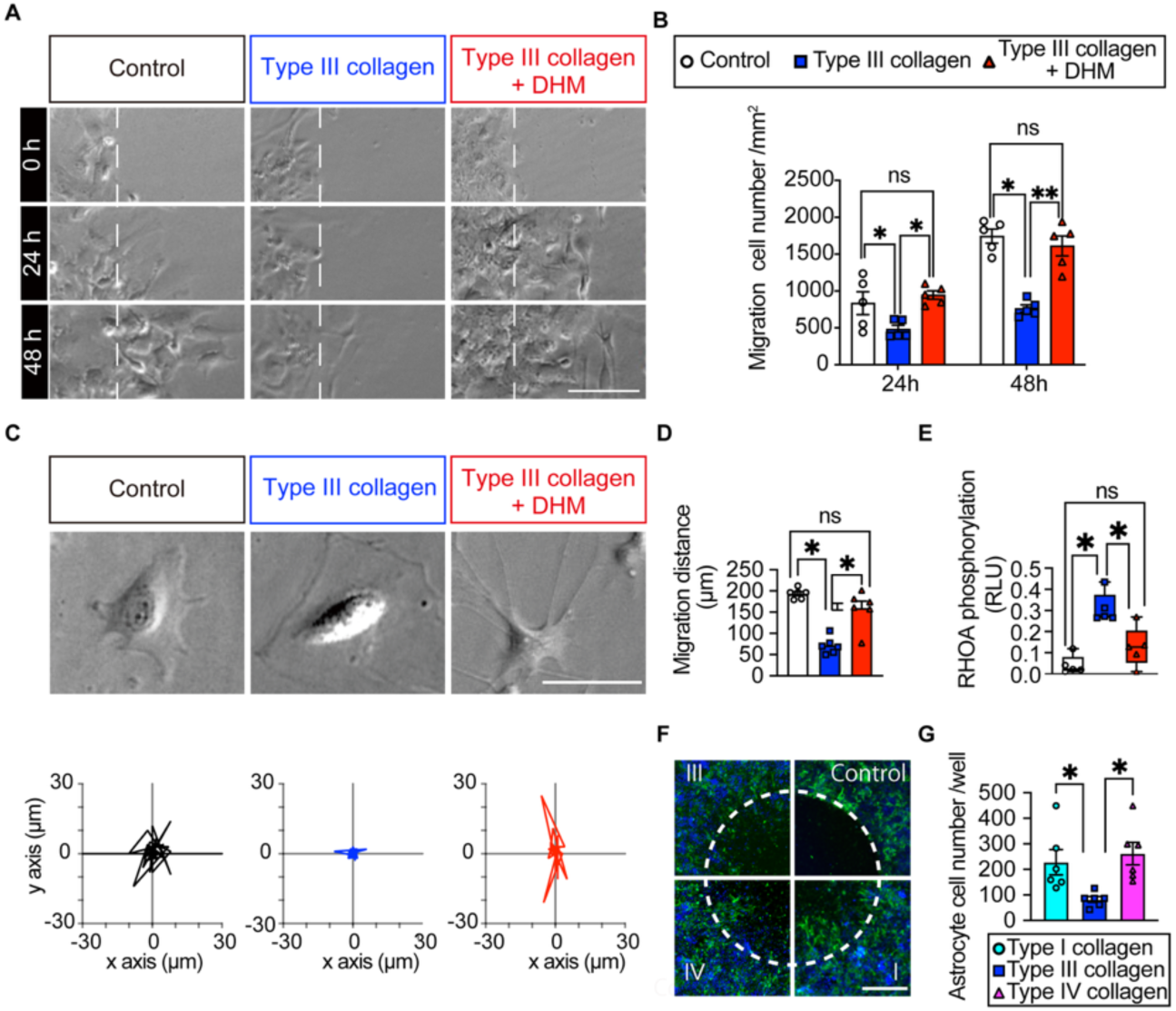
ADGRG1-type III collagen pathway controls astrocytic migration. (A) In vitro scratch injury of monolayer-cultured astrocytes on poly-L-lysine coating, type III collagen coating, or type III collagen coating with DHM. Scale bar: 100 μm. (B) Quantitative analysis of the number of migrating cells among the three groups (n = 5 per group). (C) Representative images and migration plots of astrocytes from cell tracking assay among the three groups. Scale bar: 50 μm. (D) Quantitative analysis of migration distance among the three groups. (E) Quantitative analysis of RHOA-GTP binding levels among the three groups. (F) In vitro cell migration assay of monolayer-cultured astrocytes on type III, type I, or type IV collagen coating. Control: astrocytes cultured on type III collagen with the stopper kept until fixation. Representative images depicting GFAP (green) and Hoechst (blue). Scale bar: 500 μm. (G) Quantitative analysis of the number of migrating cells among the three groups. *P < 0.05, **P < 0.005, one-way ANOVA (B, D, E, G). Data represent the mean ± s.e.m (B, D, G). Data represent minimum to maximum (E).

### Potential therapeutic efficacy of ADGRG1 antagonism in human SCI

Human-induced pluripotent stem cells (hiPSCs) have been increasingly utilized in drug screening applications in recent years (Silva & Haggarty, 2020). Using astrocytes derived from hiPSCs, we evaluated the potential of ADGRG1 blockade for human clinical application. These hiPSC-derived astrocytes expressed GFAP and ADGRG1 (Fig. 4 A and B). We divided the hiPSC-derived astrocytes into three groups: hiPSC-derived astrocytes cultured on poly-L-lysine coating (control group), type III collagen coating (type III collagen group), and type III collagen coating with DHM treatment (DHM group). The migration ability of the hiPSC-derived astrocytes was assessed using scratch assays and cell migration tracking. It was observed that the migration ability was suppressed by type III collagen and importantly, DHM treatment successfully restored their migration ability (Fig. 4 C-F). These findings indicate that the type III collagen-ADGRG1 pathway affects cell migration in human astrocytes, highlighting its potential therapeutic applications in human SCI.

**Figure 4.**
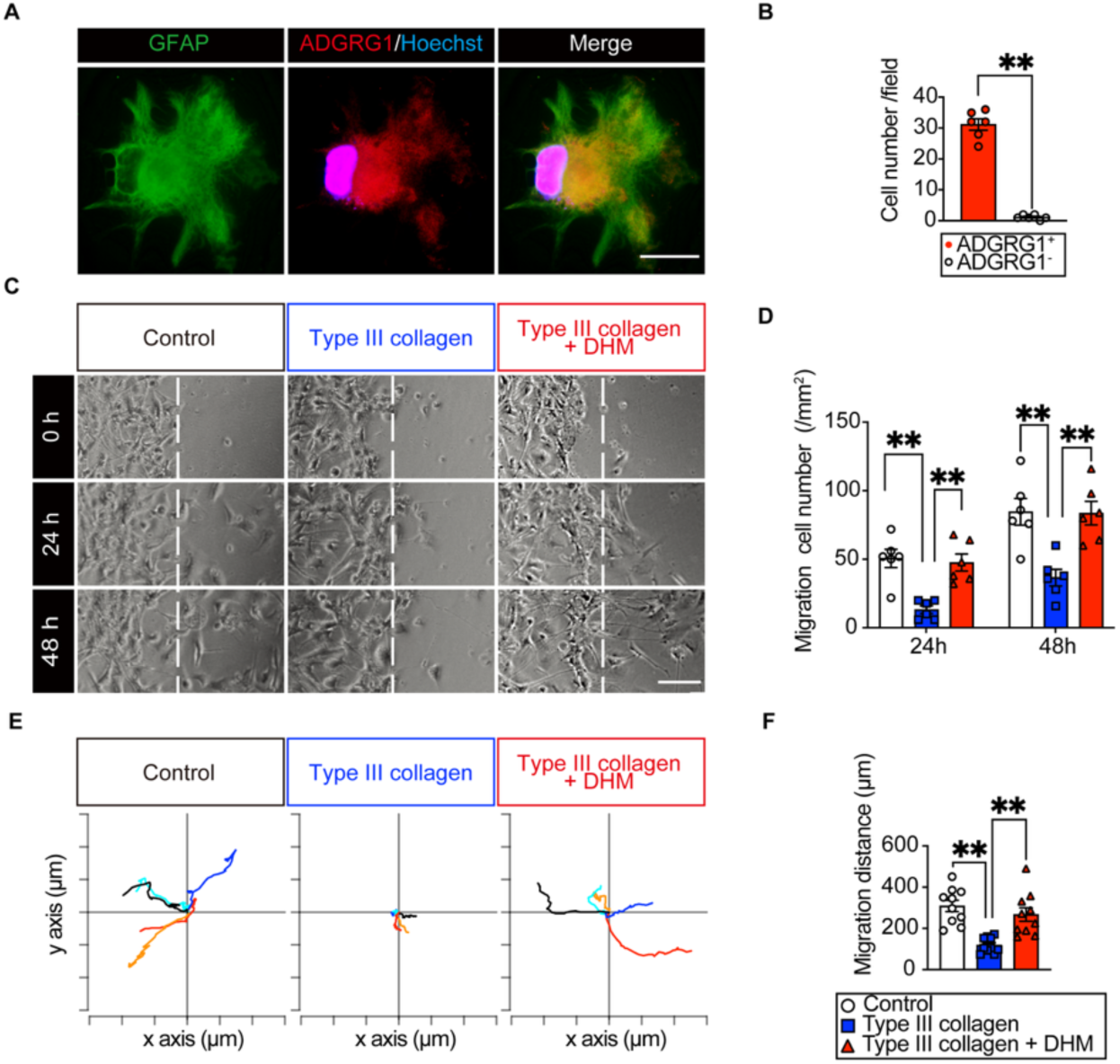
ADGRG1 blockade is effective against hiPSC-derived astrocytes. (A) Representative images of hiPSC-derived astrocytes showing GFAP (green), ADGRG1 (red), and Hoechst (blue) staining. Scale bar: 20 μm. (B) Quantitative analysis of ADGRG1-positive hiPSC-derived astrocytes. (C) In vitro scratch injury of monolayer-cultured hiPSC-derived astrocytes on Geltrex coating, type III collagen coating, or type III collagen coating with DHM. Scale bar: 500 μm. (D) Quantitative analysis of the number of migrating cells among the three groups (n = 6 per group). (E) Migration plots from the cell tracking assay of hiPSC-derived astrocytes (n = 5 per group). The tick interval is 100 μm. (F) Quantitative analysis of the migration distance among the three groups. *P < 0.05, **P < 0.005, unpaired t-test (B), one-way ANOVA (D, F). Data represent the mean ± s.e.m (B, D, F).

## Discussion

In this study, we utilized single-nucleus RNA sequencing, pathological analysis, and time-lapse imaging of astrocytes in order to elucidate the mechanisms of glial scar formation. We found that type III collagen suppressed astrocytic migration through the ADGRG1–RHOA pathway, leading to glial scar formation around the fibrotic ECM. Inhibition of ADGRG1 allowed astrocytes to enter the fibrotic ECM, resulting in the formation of smaller glial scars in the fibrotic ECM and better motor function recovery. These results indicate that type III collagen plays akey role in regulating the spatial localization of glial scars. ADGRG1 has the potential to be a therapeutic target for SCI for several reasons.

First, Blockade of the type III collagen–ADGRG1 pathway to induce glial scar formation inside the fibrotic ECM may be beneficial, considering the recently recognized dual role of glial scars. While glial scars can serve as physical barriers to axon regeneration, glial scars also have a beneficial effect on axonal growth by sealing inflammatory cells within the glial scar (Okada *et al*, 2006b). Therefore, simply removing glial scars is not necessarily beneficial (Anderson *et al*, 2016). ADGRG1 inhibition may be advantageous in that it reduces the area enclosed by the glial scar while preserving the glial scar itself. Indeed, in the DHM inj group, inflammatory responses were similar to controls, suggesting that the barrier against inflammatory cells remained intact.

Second, ADGRG1 inhibition seems to promote axon regeneration by changing the distribution of reactive astrocytes. In the DHM inj group, compact glial scars were formed more internally within the fibrotic ECM, thereby creating spaces instead of glial scars at the lesion site. These spaces appeared to be filled by reactive astrocytes, which may have contributed to creating a more permissive environment for axonal growth. Consistent with this notion, corticospinal tract (CST) fibers in the DHM inj group frequently projected across the lesion site and, on the caudal side, often took alternative routes rather than passing through the dorsal column (Fig. 2E). Since CST axons in mice typically traverse the dorsal column (Moreno-Lopez *et al*, 2021), this observation suggests the formation of new detour circuits rather than mere sparing of pre-existing fibers, supporting true axonal regeneration. Notably, the therapeutic effect was strongest when DHM was administered at 4 dpi (Fig. S2 A–E), a time point when Astro_4 cells were the dominant subtype (Fig. 1B). Astro_4 cells are characterized by high expression of *Spp1* (Fig. S1B), a gene encoding a glycoprotein that promotes neurite outgrowth (Li & Jakobs, 2022). Thus, the redistribution of *Spp1*-expressing Astro_4 cells into the newly generated scar-free areas may underlie the enhanced and more efficient axonal regeneration observed in our study.

Finally, our study identified type III collagen–ADGRG1 interaction as a critical upstream event that drives excessive activation of RHOA, thereby retaining astrocytes around the fibrotic ECM. Because both excessive and insufficient RHOA activity can impair cell migration (Renault-Mihara & Okano, 2017; Li *et al*, 2018; Ray *et al*, 2002), direct manipulation of RHOA may be suboptimal. Targeting the upstream cause of excessive activation—binding of type III collagen to ADGRG1—may therefore be a more effective and physiologically balanced strategy than conventional approaches aimed at RHOA itself. In conclusion, type III collagen traps astrocytes around fibrotic ECM via the astrocytic ADGRG1–RHOA pathway. ADGRG1 blockade shifts glial scar formation from the edge of the fibrotic ECM into its interior, reducing scar volume and facilitating axonal regeneration. Our findings provide a deeper insight into the role of type III collagen in forming glial scars, demonstrating the potential therapeutic application of blockade of astrocytic ADGRG1 after traumatic CNS injury.

## Materials and Methods

### Structured Methods - Reagents and Tools Table

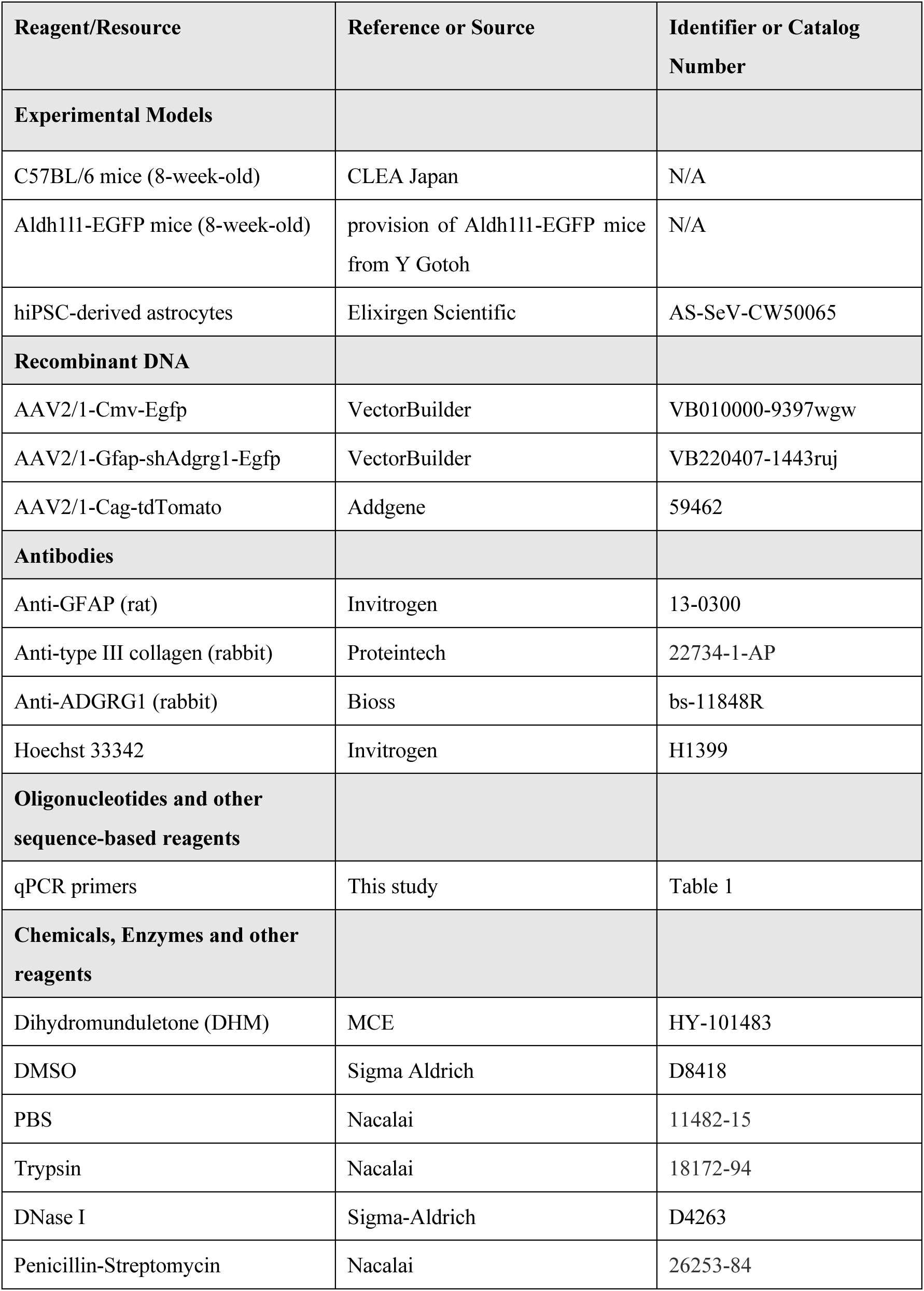
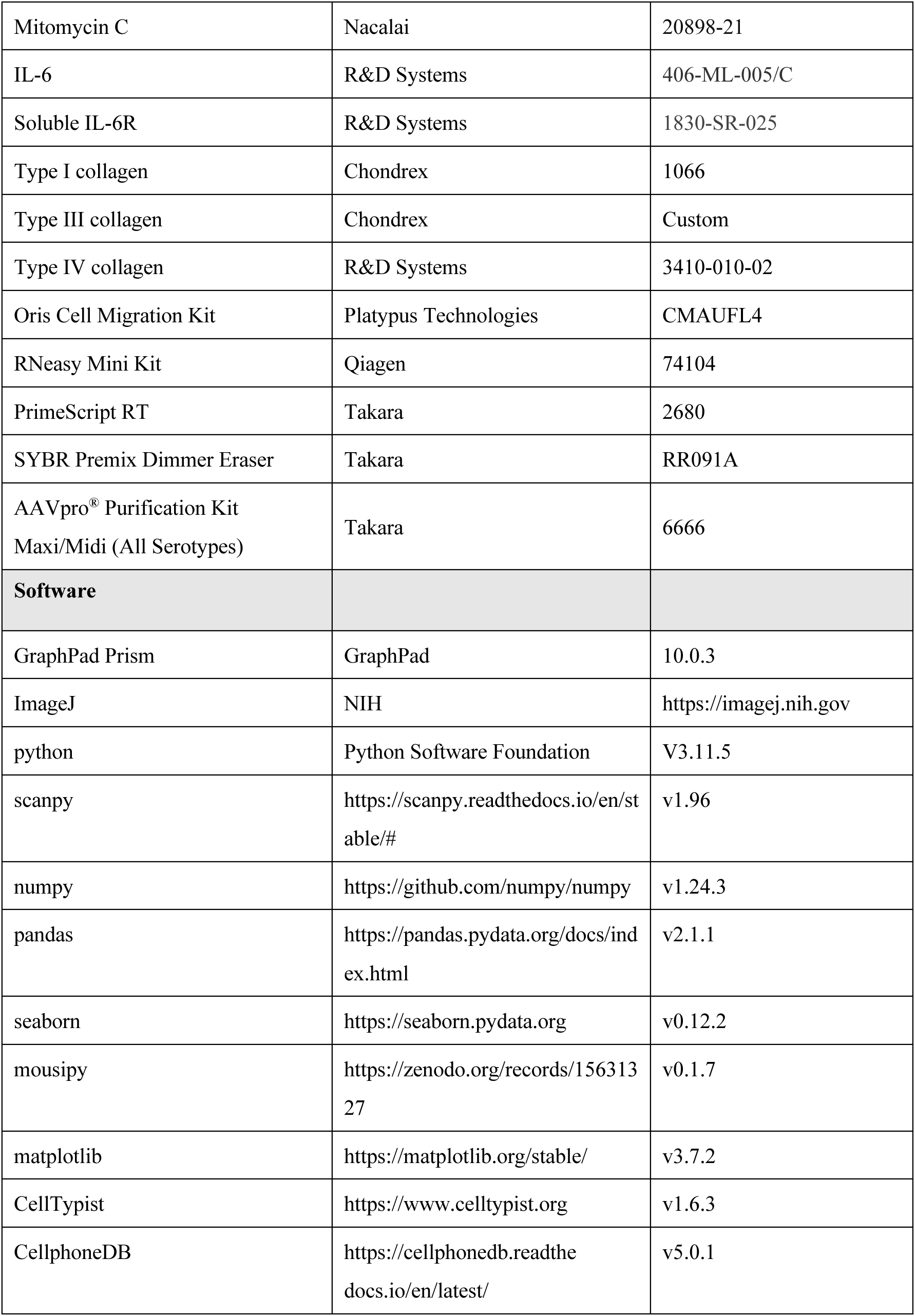

### Animals

All mice were housed in a temperature- and humidity-controlled environment with a 12-hour light-dark cycle and *ad libitum* access to food and water. C57BL/6 mice (8-week-old) and Aldh1l1-EGFP mice (8-week-old) were used in this study.

### Materials

DHM (HY-101483; MCE) (Stoveken *et al*, 2016b) was dissolved in DMSO to a final concentration of 1 mM to prepare a stock solution. For cell culture experiments, the stock solution was diluted with culture medium to the desired concentration. For animal administration experiments, it was diluted with PBS to the appropriate concentration and used accordingly.

### Spinal Cord Injury

The mice were anesthetized with 2% isoflurane and subjected to contusion injury (70 kilodynes) at the 10th thoracic level using an Infinite Horizons Impactor (Precision Systems Instrumentation) (Kobayakawa *et al*, 2014). After the injury, the overlying muscles were sutured, and the skin was closed with wound clips. During the period of recovery from anesthesia, the animals were placed in a temperature-controlled chamber until thermoregulation was reestablished. The motor function was evaluated using the locomotor open-field rating scale (BMS) (Ma *et al*, 2001).

### Drug Injection

A glass tip was inserted into the epicenter of the injured spinal cord, and 2 μl of 100 μM DHM (HY-101483; MCE) was injected at 0.5 μl/min using a stereotaxic injector (KDS 310; Muromachi Kikai) at 4 dpi (Kobayakawa *et al*, 2019). Control animals received 2 μl of 0.1 M PBS at 4 dpi.

### Primary Astrocyte Cultures

Purified primary astrocyte cultures were prepared from C57BL/6 mice, as previously described (Schildge *et al*, 2013; Okada *et al*, 2006a). In brief, after removal of the meninges, on postnatal day 2, mouse brain tissues were minced and incubated in a water bath at 37°C for 30 min in DMEM (08456-36; Nacalai) in the presence of 0.25% trypsin (35554-64; Nacalai) and 300 g/ml DNase I (Sigma-Aldrich). Dissociated cells were triturated with 0.25% FBS and centrifuged at 300 ×*g* for 3 min. Following dilution with astrocyte-specific medium (DMEM containing 10% FBS and 1% penicillin-streptomycin), the cells were plated on a poly-L-lysine-coated T75 flask. After 7 days in a humidified CO_2_ incubator at 37°C, a T75 flask was set up on an orbital shaker to remove microglia at 180 rpm for 30 minutes. Fresh astrocyte culture medium (20 mL) was added, and the flask was shaken at 240 rpm for 6 hours to remove the oligodendrocyte precursor cells. Astrocytes were detached from T75 flasks using 0.25% trypsin. Astrocytes were stimulated with 50 ng/ml IL-6 (R&D Systems) and 200 ng/ml soluble IL-6 receptor (R&D Systems) for one day. Regular mycoplasma testing was performed, and all results were negative throughout the course of the present experiments. Testing was performed using MycoStrip (rep-mys-10; INVIVOG).

### hiPSC-derived Astrocytes Cultures

We used hiPSC-derived astrocytes purchased from Elixirgen Scientific (AS-SeV-CW50065; Elixirgen Scientific). The cells were cultured for 14 days, as specified by the kit, and used for the experiments.

### Scratch Assay

Mouse astrocytes were plated on coverslips coated with poly-L-lysine or type III collagen. After reaching sub-confluency, we treated cells with 10 mg/ml mitomycin C for 2 h to avoid the effects of cell proliferation and then scratched them. Similarly, hiPSC-derived astrocytes were plated on coverslips coated with Geltrex (A1569601; Thermo Fisher Scientific) or type III collagen (PSC-3-100-05; Nippi). In DHM group, we added DHM to the medium to a final concentration of 20 μM. We created a cell-free area by scratching the monolayer with a pipette tip and evaluated the migration of cells to the cell-free area from the surrounding area for 48 hours. After capturing photographs of five (mouse astrocytes) or six (hiPSC-derived astrocytes) non-overlapping fields, we counted the number of migrating astrocytes.

### Cell Tracking Assay

Mouse astrocytes were seeded at a low density (approximately 30% confluency) with poly-L-lysine or type III collagen. For hiPSC-derived astrocytes, cells were plated at low density on coverslips coated with Geltrex or type III collagen. To assess dynamic astrocyte behavior, time-lapse imaging was performed under a phase-contrast microscope equipped with a temperature- and CO₂-controlled incubator. Images of six (mouse astrocytes) or ten (hiPSC-derived astrocytes) astrocytes per condition were acquired every 10 minutes for a total of 6 hours. For the DHM-treated group, DHM was added to the culture medium at a final concentration of 20 μM prior to initiating time-lapse imaging. The resulting image sequences were analyzed to track individual astrocyte migration and quantify morphological changes using ImageJ (NIH). Key parameters included cell migration distance, speed, and process length.

### Immunohistochemistry (IHC)

Mice were anesthetized and transcardially perfused with normal saline, followed by 4% paraformaldehyde (PFA) in 0.1 M phosphate-buffered saline. The spinal cord was removed and immersed in 4% PFA at 4°C for 24 hours. A spinal segment centered over the lesion epicenter was transferred into 10% sucrose in PBS for 24 hours and 30% sucrose in PBS for 24 hours and embedded in O.C.T. compound. The embedded tissue was immediately frozen in liquid nitrogen and stored at -80°C until use. Frozen sections were cut with a cryostat in the sagittal or axial plane at 16 μm and mounted onto glass slides as previously described (Kijima *et al*, 2019). For immunofluorescence staining, the spinal cord sections were permeabilized with 0.01% Triton X-100 and 10% normal goat serum in PBS (pH 7.4) for 60 minutes. The sections were then stained with primary antibodies against GFAP (1:500; astrocyte marker, rat; 13-0300; Invitrogen), Type III collagen (1:500; rabbit; Protein Tech), or ADGRG1 (1:500; rabbit; bs-11848R; Bioss). The sections were then incubated with Alexa Fluor-conjugated secondary antibodies (1:1000; Invitrogen). Nuclear counterstaining was performed using Hoechst 33342 (1:1000; Invitrogen). All images were captured using a BZ-X700 digital microscope system (Keyence) or epifluorescence microscope equipped with a digital camera (BX51; Olympus). Scar volume was quantitatively evaluated by measuring the scar area in every serial section taken at 80 μm intervals and approximating the scar volume. GFAP-positive area inside type III collagen, the section showing the maximum scar area was used, and the GFAP-positive area within the region of dense type III collagen staining was measured. All measurements were performed using ImageJ.

### Cell Migration Assay

Cells (5 × 10^4^) were seeded into wells of the Oris Cell Migration Assembly Kit-FLEX (CMAUFL4; PlatypusTechnologies), coating with type I collagen (1066; Chondrex), type III collagen (custom order; Chondrex), or type IV collagen (3410-010-02; R&D Systems). Migration assays were conducted in accordance with the manufacturer’s instructions. After attached for 16 hours, cells were allowed to migrate into the clear field after removal of the well inserts. Cells were then fixed with 4% PFA and stained with primary antibodies against GFAP. Nuclear counterstaining was performed using Hoechst 33342. The images were analyzed using the ImageJ.

### Quantitative Real-time Polymerase Chain Reaction (qPCR)

Total RNA was isolated from astrocytes and spinal cord tissues using a RNeasy Mini Kit (74104; Qiagen). cDNA was synthesized from total RNA using PrimeScript Reverse Transcriptase (2680; Takara), according to the manufacturer’s instructions. RT-qPCR was performed using primers specific to the genes of interest and SYBR Premix Dimmer Eraser (RR091A; Takara). The data were normalized to glyceraldehyde-3-phosphate dehydrogenase (GAPDH) levels. RT-qPCR was performed using a CFX Connect Real-Time PCR Detection System (Bio-Rad). Using one cDNA sample, we examined the mRNA expression of the various factors listed in Table 1.

**Table 1.**
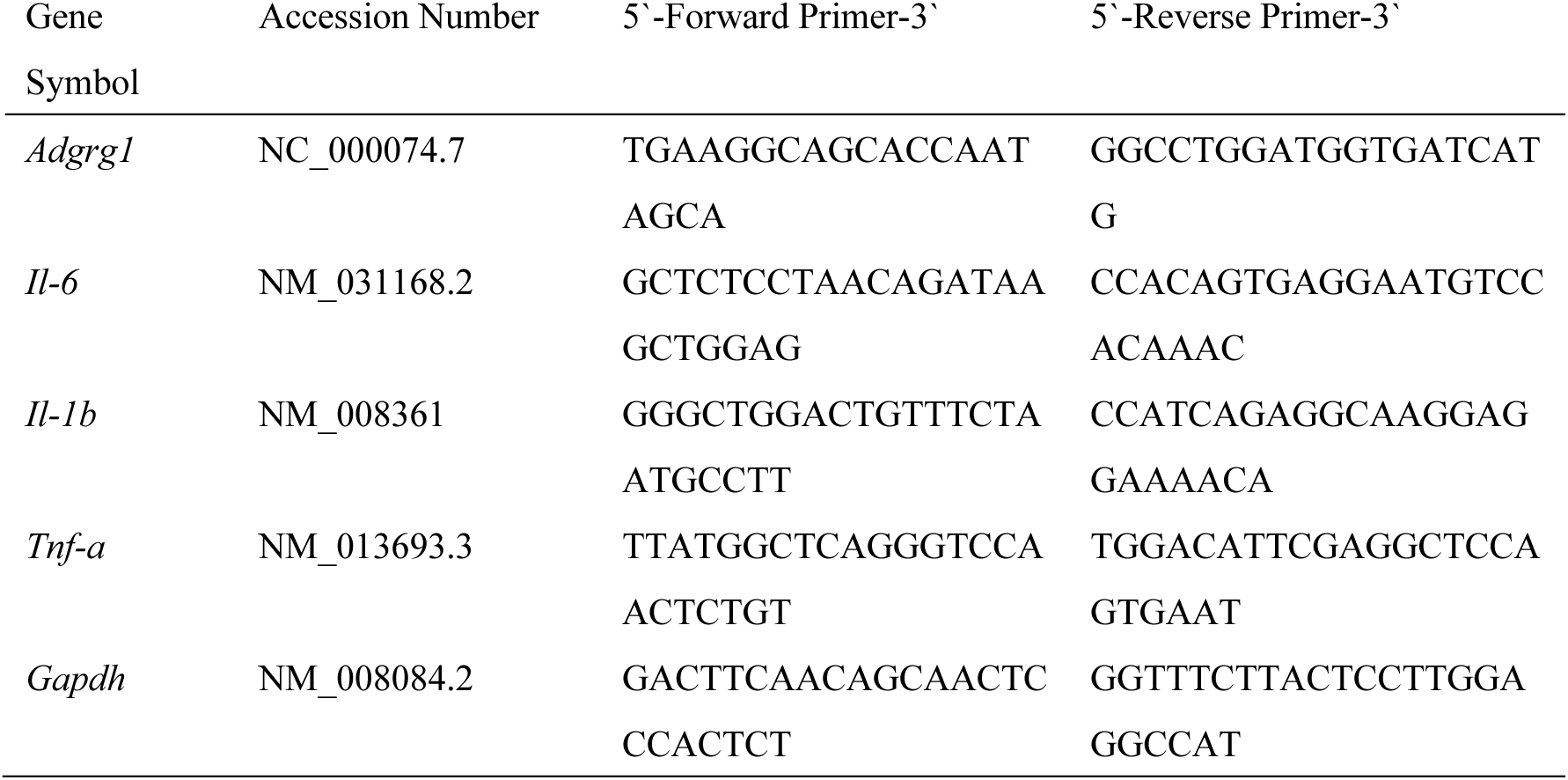
Primers used for quantitative RT-PCR.

### Analyses of the Locomotor Function

The motor function of the paralyzed hind paws was evaluated using a locomotor open-field rating scale on the Basso Mouse Scale (BMS) and a footprint analysis. For BMS, each mouse was assessed at 1, 4, 7, 14, 21, 28, 35 and 42 dpi. A team of three independent examiners evaluated each animal for 4 minutes and assigned an BMS score. The median score of the three examiners was used for analysis. All tests were performed in a double-blinded manner as previously described (Kobayakawa *et al*, 2014).

### AAVs

Various adeno-associated viral vectors (AAV) were used to deliver either control AAV (AAV2/1-*Cmv-Egfp*; 1 × 10^13^ vg/ml; VB010000-9397wgw; VectorBuilder), *Adgrg1* knockdown (AAV2/1-*Gfap-shAdgrg1-Egfp*; 1 × 10^13^ vg/ml; VB220407-1443ruj; VectorBuilder), or AAV-*Cag-tdTomato* (AAV2/1-*Cag-tdTomato*; 1 × 10^13^ vg/ml; 59462; Addgene). A glass tip was inserted into the epicenter of the spinal cord for *Adgrg1* knockdown or the hemisphere for AAV-*Cag-tdTomato*, and AAV was injected at 0.5 μl/min using a stereotaxic injector (KDS 310; Muromachi Kikai).

### Neuronal Tracing

At 14 dpi, a craniotomy was performed to expose the sensorimotor cortex. tdTomato-encoding AAV was injected into the hemisphere. Six mice per group received AAV-*Cag-tdTomato*. For the mouse, three injections for cortex (AAVs; 1.0 × 10^13^ virus genome/mL, 1 μL per site) were performed at the following coordinates: 1.0 mm lateral, 1.0 mm depth; and -1.0 mm, 0 mm, 1.0 mm posterior to the bregma. Six weeks post-SCI (4 weeks post-AAV injection), the spinal cord was fixed with PFA, as described above. Sections were taken on the caudal and rostral sides (1.5 mm) from the center of the injury to assess how tdTomato-positive areas projected to the caudal side in comparison to sections on the rostral side (Hirota *et al*, 2022).

### Three-dimensional Reconstruction of Immunostaining

Spinal cord sequential sections at 80-μm intervals were stained with antibodies against GFAP and type III collagen; type III collagen-positive areas surrounded by strongly GFAP-positive areas were defined as scar and non-scar tissue was removed. Stacked images were created by positioning the images with markers. The stacked images were converted to 3D data using the ImageJ software program.

### Flow Cytometry

Spinal cords (4.0 mm in length, centered around the lesion) were dissociated using collagenase type I (17100017; Invitrogen), as previously described (Saiwai *et al*, 2010). The ALDH1L1-positive cells or AAV-infected astrocytes were sorted as EGFP-positive cells using a FACSAria Fusion Flow Cytometer and the FACSDiva software program (BD Biosciences).

### Single Nuclear RNA Sequencing (snRNA-seq) Analysis

We analyzed public snRNA-seq datasets with Scanpy v.1.7.0 (Wolf *et al*, 2018), with the pipeline following their documentation. Specifically, clustering was performed using the Leiden algorithm. Cell annotation was performed with Cell Typist with modified name for spinal cord tissue (Domínguez Conde *et al*, 2022; Xu *et al*, 2023; Yao *et al*, 2023). Cell interaction analysis was performed using Cellphone DB v5.0 (Garcia-Alonso *et al*, 2022), with the pipeline after translating with mousipy.

### Spatial Transcriptomics

We analyzed public spatial transcriptomics datasets using Scanpy v.1.7.0 (Wolf *et al*, 2018). As a reference, we utilized the snRNA-seq dataset and performed supervised label transfer to map predicted cell type annotations onto the spatial transcriptomics data. Specifically, cell types identified in the snRNA-seq data were transferred to the spatial transcriptomics dataset by matching shared marker gene expression profiles and predicted labels. We then integrated the normalized gene expression matrices with spatial coordinates to construct 3D spatial representations. To explore local spatial interactions, we applied K-nearest neighbors classification (n = 5) to the coordinate data, identifying adjacent cells and calculating the average expression of target genes in their neighborhood. We selectively visualized the astrocytes located near *Col3a1*-expressing cells using matplotlib, emphasizing the spatial distribution and microenvironmental interactions of the annotated cell types.

### Statical Analysis

All statistical analyses were performed using GraphPad Prism 10 version 10.0.3 (GraphPad Software Inc.) following the suggested test parameters. Details regarding the statistical analysis of experiments throughout the study can be found in the figure legends.

## Data availability

Raw experimental data generated in this study are provided in the source data accompanying this paper. For the RNA sequencing analyses, publicly available data from the Gene Expression Omnibus (GEO) under accession number GSE234774 were utilized. All codes and data necessary to reproduce the findings of this study are included in the source data. Any additional information required for reanalysis is available from the corresponding author upon reasonable request.

## Author contributions

Gentaro Ono performed the experiments and analyzed the data. Kazu Kobayakawa, Hirokazu Saiwai, and Kazuya Yokota supervised the research design. Tetsuya Tamaru, Kazuhiro Hata, Hirotaka Iura, Yohei Haruta, Kazuki Kitade, Jun Kishikawa, Kiyoshi Tarukado, Kenichi Kawaguchi, Takeshi Maeda, Seiji Okada, and Yasuharu Nakashima contributed equally to this study. Dai-Jiro Konno and Kinichi Nakashima provided guidance on the experimental technology. Gentaro Ono wrote the original draft. Kazu Kobayakawa, Hirokazu Saiwai, and Kazuya Yokota reviewed and edited the manuscript. Kazu Kobayakawa designed the studies, supervised the overall project, and prepared the final version of the manuscript. All authors have read and approved the final manuscript.

## Competing interests

The authors declare no competing interests.

## Acknowledgements

We thank Y. Gotoh for providing the Aldh1l1-EGFP mice. We also appreciate the technical assistance from the Research Support Center, Research Center for Human Disease Modeling, Kyushu University Graduate School of Medical Sciences. This study was supported by JSPS KAKENHI Grant Numbers (18K16665, 19K18515, 22K09426, 22K19587, 23K27722, and 23H03031), JST FOREST Program (JPMJFR220P), ZENKYOREN (National Mutual Insurance Federation of Agricultural Cooperatives), Takeda Science Foundation, and the General Insurance Association of Japan.

---

**Figure S1.**
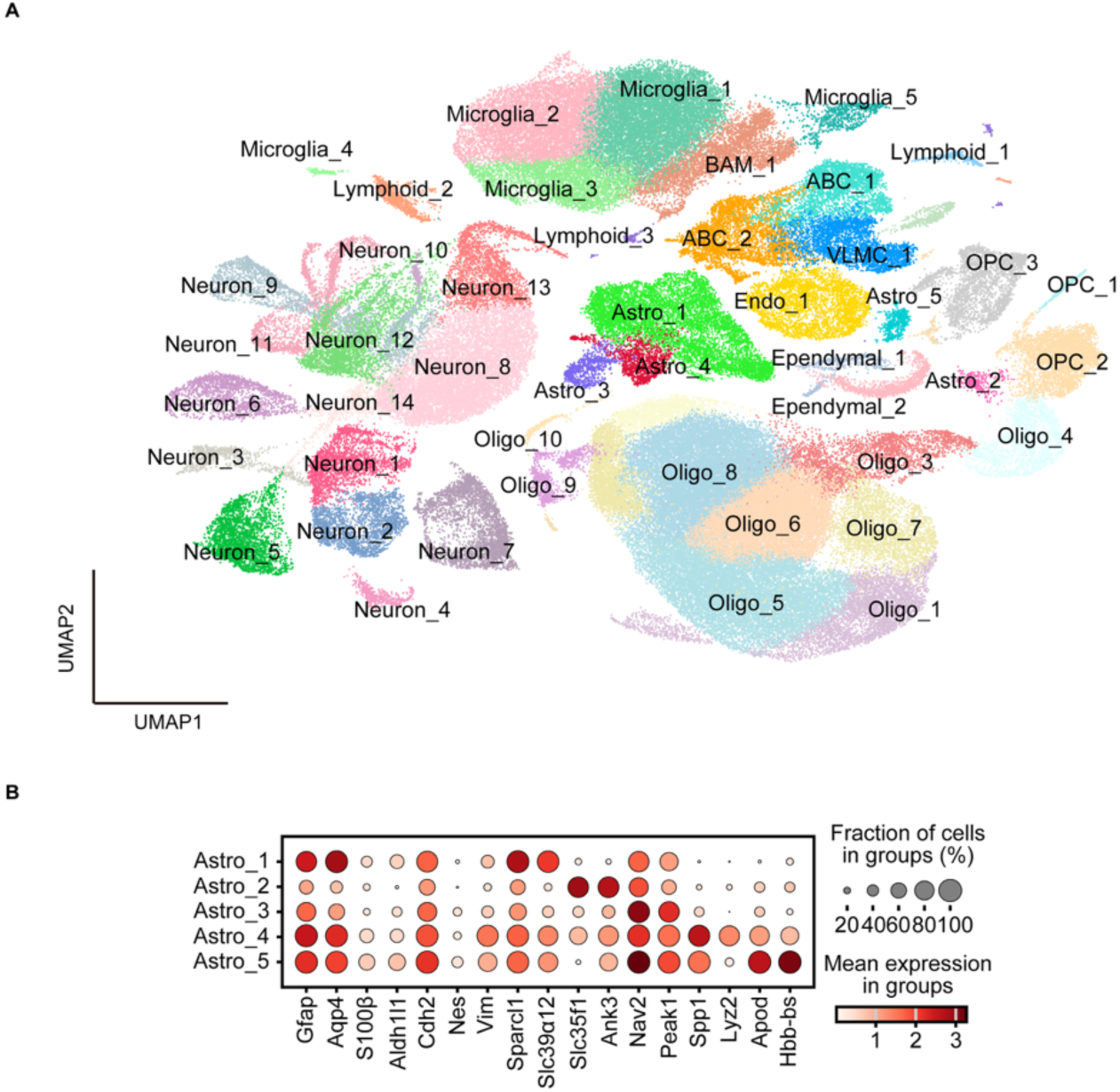
snRNA-seq analysis overview. (A) Detailed UMAP of cell states (cell types) in the injured spinal cord snRNA-seq (n = 151712) datasets. Astro: Astrocytes, Oligo: Oligodendrocytes, Microglia, Neuron, BAM: Border-associated macrophages, OPC: Oligodendrocyte precursor cells, ABC: Arachnoid barrier cells, Endo: Endothelial cells, VLMC: vascular leptomeningeal cells, Ependymal: Ependymal cells, Lymphoid: Lymphoid cells, Peri: Pericytes. (B) Dot plots showing the variance-scaled expression of marker genes in astrocytes.

**Figure S2.**
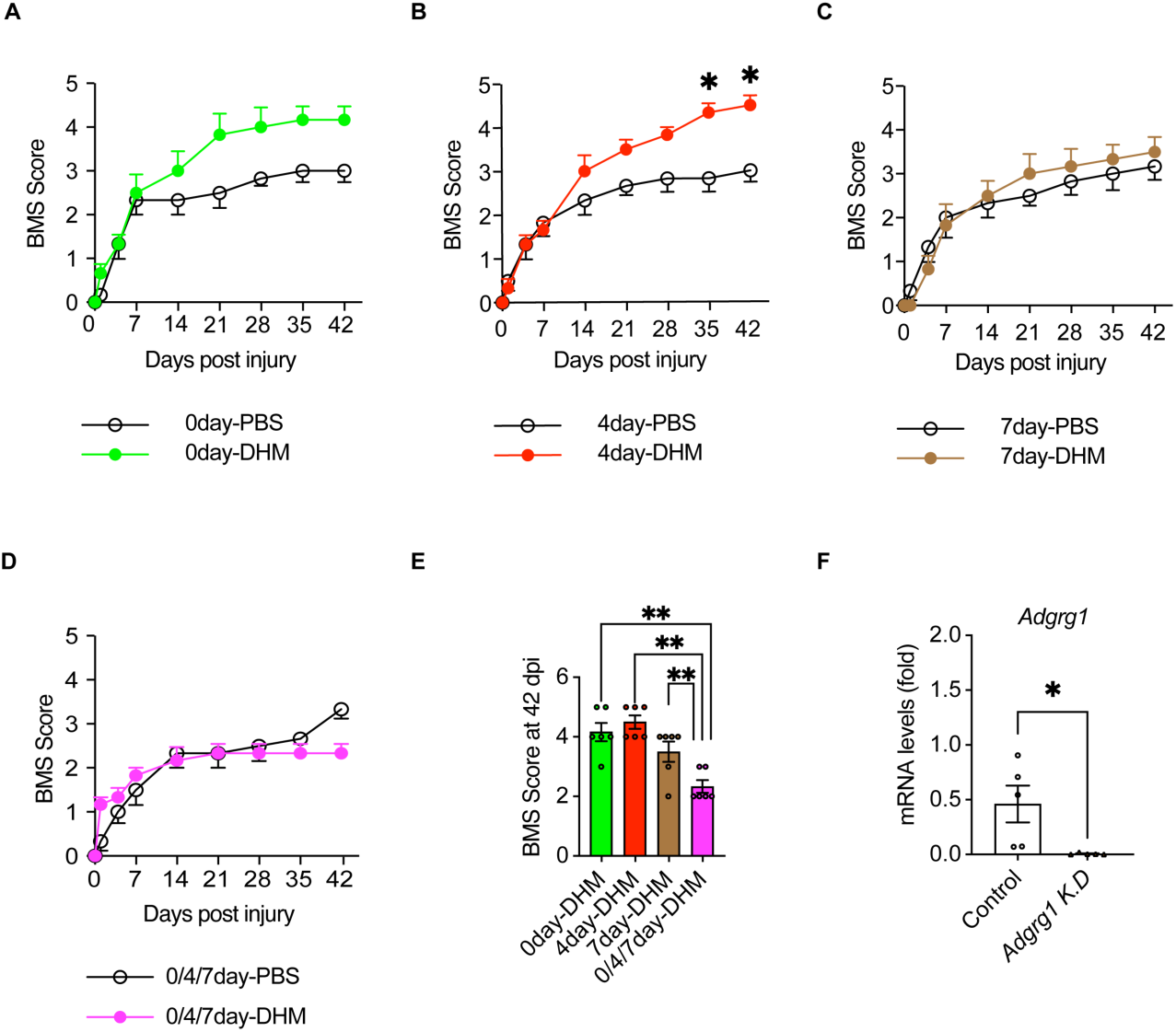
The efficacy of ADGRG1 inhibition depends on the time point of intervention. (A) Time course of the BMS score after SCI of the groups receiving PBS injection and DHM injection at 0 dpi (n = 6 mice per group). (B) Time course of the BMS score after SCI of the groups receiving PBS injection and DHM injection at 4 dpi (n = 6 mice per group). (C) Time course of the BMS score after SCI between the groups receiving PBS injection and DHM injection at 7 dpi (n = 6 mice per group). (D) Time course of the BMS score after SCI of the groups receiving PBS injection and DHM injection at 0, 4, and 7 dpi (n = 6 mice per group). (E) BMS score at 42 dpi between the groups receiving DHM injection (n = 6 mice per group). (F) *Adgrg1* expression between naïve astrocytes and astrocytes infected with AAV-*Gfap*-*shAdgrg1*-*Egfp.* *P < 0.05, **P < 0.005, two-way ANOVA (A-D), one-way ANOVA (E), unpaired t-test (F). Data represent the mean ± s.e.m (A-F).

**Figure S3.**
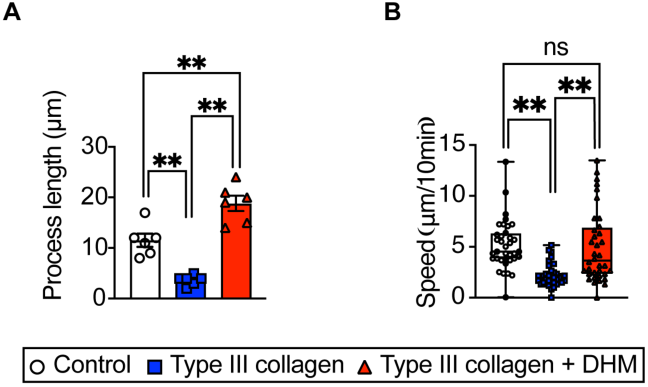
Detailed analysis of astrocyte migration. (A) Quantitative analysis of astrocyte process length cultured on poly-L-lysine, type III collagen, or type III collagen with DHM. (B) Quantitative analysis of migration speed among the three groups.

## Notes

### Competing Interest Statement

The authors have declared no competing interest.

